# Open prediction of polysaccharide utilisation loci (PUL) in 5414 public *Bacteroidetes* genomes using PULpy

**DOI:** 10.1101/421024

**Authors:** Rob D. Stewart, Marc D. Auffret, Rainer Roehe, Mick Watson

## Abstract

Polysaccharide utilisation loci (PUL) are regions within bacterial genomes that encode all the necessary machinery for the cleavage of particular carbohydrates. For the *Bacteroidetes* phylum, prediction of PUL from genomic data alone involves the identification of carbohydrate-active enzymes (CAZymes) co-localised with *susCD* gene pairs. Here we present the open prediction of PUL in 5414 public *Bacteroidetes* genomes, and an open-source pipeline to reproduce or extend the results.

## Background & Summary

Glycans are ubiquitous and abundant in living organisms, include important molecules such as glycogen, chitin, cellulose, hemicellulose and pectin, and have roles in cell structure, energy storage and signalling. They also form the most important carbon source for the majority of organisms on Earth, with many microorganisms encoding a diverse array of carbohydrate active enzymes (CAZymes) to aid the breakdown of complex carbohydrates into simpler sugars.

In ruminants such as cattle, buffalo, goats and sheep it is largely the rumen microbiota which are responsible for the breakdown of recalcitrant complex carbohydrates present in fibrous plant diets into more simple sugars and short-chain fatty-acids that the host can use for production of meat and milk^1^. Unsurprisingly, metagenomic sequencing of the rumen microbiota has revealed a large number of CAZymes focused on the digestion of an array of carbohydrate substrates^2–8^. The human diet is also abundant in complex dietary carbohydrates, and similar efforts to characterise CAZymes in the human gut exist^9–11^.

Members of the *Bacteroidetes* phylum have been found in almost all environments studied to date and are particularly abundant in animal digestive systems. Many members of the *Bacteroidetes* encode polysaccharide utilisation loci (PUL) in their genomes. A PUL is a genomic locus that encodes all the necessary machinery to bind a particular class of polysaccharide at the cell surface, perform an initial cleavage and import the products inside the cell. Characterised PUL in *Bacteroidetes* generally consist of a *susC-susD* pair with a number of CAZymes encoded close by^12,13^. The *SusCD* proteins are responsible for binding the carbohydrate, and the CAZymes for cleavage.

Many genome annotation tools exist^14^, but certain classes of protein/enzyme require more in-depth analysis, such as carbohydrate-active enzymes (CAZymes), often a focus of bioprospecting by the industrial biotechnology community^15^. PULDB (http://www.cazy.org/PULDB/) and CAZy (http://www.cazy.org/) are the two most important databases for PUL and CAZyme annotation respectively. The CAZy database is the “industry standard” database for carbohydrate active enzymes and represents the most comprehensive resource yet published. CAZy is updated every few weeks and includes both automatic and manual annotation^16^. PULDB is built upon CAZy – initially published as an algorithm to predict PUL in 67 *Bacteroidetes* genomes^12^, a recent update extended this to 820 genomes^13^. Like CAZy, PULDB consists of automatic predictions and manually curated information.

Despite their immensely important status as “industry standard” databases, both CAZy and PULDB suffer from problems. As others have noted^17,18^, CAZy and PULDB are optimised for simple, single user queries. Bulk querying, bulk download and bulk search are not enabled; annotation of new or user genomes is not possible; and certain aspects of the annotation methods are opaque. Furthermore, despite the PUL prediction algorithm of PULDB having been published^12^, to our knowledge no software implementation was released.

In this manuscript we present the automated, reproducible and scalable prediction of PUL in 5414 public *Bacteroidetes* genomes. The predictions are fully open and can be accessed and used by any researcher, commercial or otherwise. The analysis workflow (which we have called “PULpy”) is also released as a Snakemake pipeline^19^, freely available on GitHub. Whilst ours are purely computational predictions, we show significant overlap with PULDB predictions with minimal false positives and negatives, and conclude that this work represents the largest and first fully open^20^ and accurate prediction of PUL in *Bacteroidetes* genomes.

## Methods

### Analysis

A flowchart of the analysis workflow can be seen in figure 1. There are two starting points to the workflow – either a genome with no annotation, or a genome with annotation in NCBI format.

**Figure 1.**
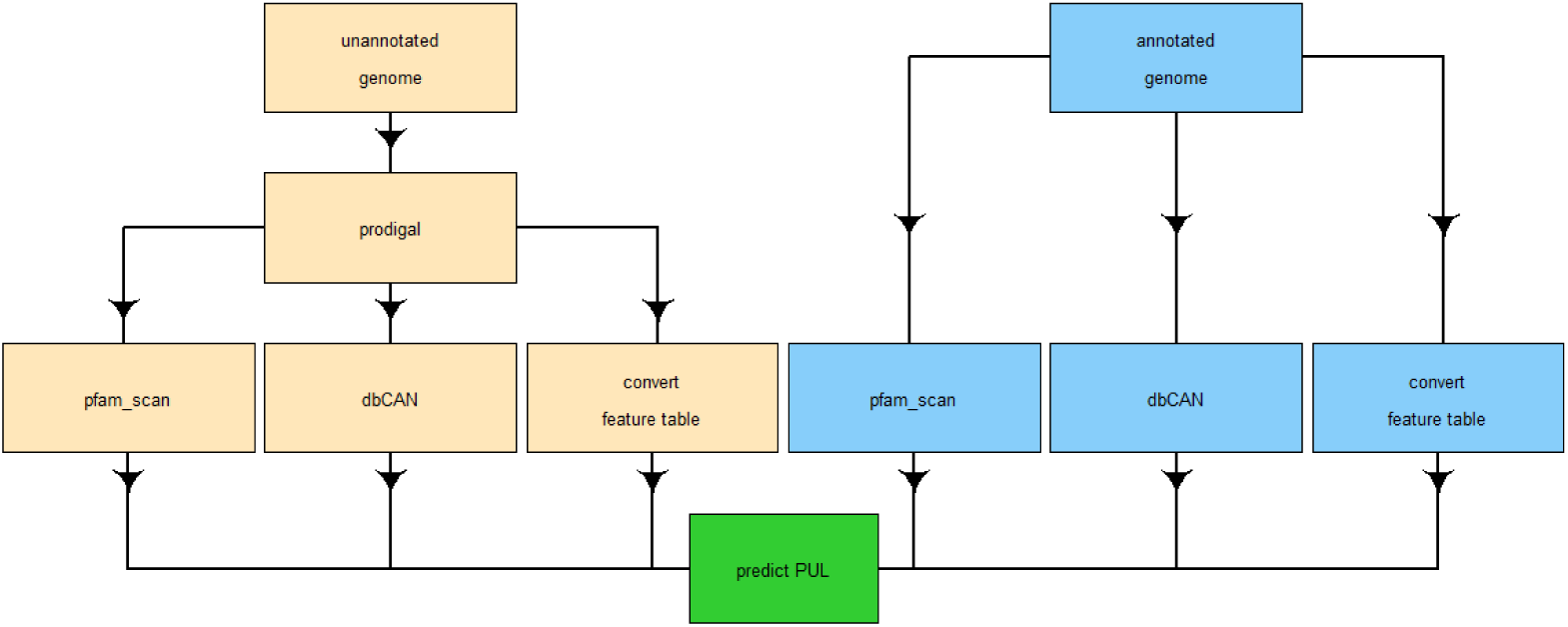
Flowchart of the steps taken in PULpy. Users may start with an unannotated genome, or with the NCBI genome annotation.

A genome with no annotation is presented to PULpy as a simple zipped FASTA file. Proteins are predicted with Prodigal^21^, outputting both a protein FASTA file and a GFF describing the location of the predicted gene and protein features. The GFF is converted to a simple standardised feature table format used by PULpy. Simultaneously, the protein FASTA file is searched against Pfam^22^ using pfam_scan.pl and dbCAN^18^ (a database of CAZy families) using HMMER3^23^. A genome with annotation is defined as one that has an existing protein FASTA file, and annotation in NCBI feature table format. If these are present, then the existing protein FASTA is used for Pfam and dbCAN searches, and the NCBI feature table format is converted to PULpy’s internal format.

The Pfam results are used to identify potential *susCD* pairs, and the dbCAN results enable the identification of the different CAZyme families – glycoside hydrolases, glycosyl transferases, polysaccharide lyases and carbohydrate esterases. The feature table allows PULpy to cross-reference *susCD* and CAZyme annotation with genomic position, and the subsequent prediction of PUL.

Prediction of PUL is carried out by an R script that reads the Pfam, dbCAN and feature table data tables. PULpy employs a sliding window approach, very similar to that described in Terrapon *et al*^12^. It is very difficult to recreate the algorithm exactly as it is not sufficiently described in the paper to enable precise reproducibility; and occasionally, the results in PULDB appear to contradict the rules described in the paper (presumably due to subsequent manual annotation). PULpy takes the following approach. First, *susCD* pairs are found. Secondly, PULpy looks downstream using a sliding window of five genes, with the caveat that the intergenic distance can be no more than 500 bp. If a CAZyme (or another *susC/susD*) is predicted within that window, the window shifts one gene downstream and repeats the process. This is repeated until no further CAZymes or *susC/D* genes are found within the window, or the intergenic distance exceeds 500 bp. The entire process is then repeated upstream of the *susCD* pair. Once the downstream and upstream searches have completed, the predicted PUL is output in two formats. Firstly, a single row summary format, describing the start, end and pattern of the PUL; secondly, a tabular format, with one row per gene within the PUL.

### Input genomes

The predictions presented here arise from 5414 public *Bacteroidetes* genomes downloaded from the NCBI FTP site on July 16^th^ 2018. The entire list of input genomes is available as Supplementary Table 1. We downloaded file “assembly_summary.txt” from ftp://ftp.ncbi.nlm.nih.gov/genomes/genbank/bacteria/ and added taxonomic information using Ete3^24^. We use the GenBank assembly accession number (typically beginning GCA_xxxx) as a consistent identifier throughout the analyses and results.

Where all of a genome FASTA, a protein FASTA and annotation in NCBI feature table format were present, these were used. Otherwise, only the genome FASTA was used.

## Code availability

All code to reproduce this analysis is available at https://github.com/WatsonLab/PULpy

## Data Records

The data are freely available in the University of Edinburgh DataShare repository under DOI: 10.7488/ds/2438

Two data tables are included relating to PUL. The first is a summary, with one row per PUL prediction, describing the location and the “pattern” of the PUL. The second is a data table containing the full results, with one row per gene. This table describes the location of each gene, and the *susCD*/CAZyme annotation.

Included are a README.txt describing the columns of the data tables, and a samples.tsv file indicating which genomes were included in the analysis.

In total we predict 96117 PUL, including 528,509 genes, on 4582 public *Bacteroidetes* genomes.

## Technical Validation

A comprehensive comparison of the PULpy predictions with PULDB is impossible as PULDB is not optimised for bulk queries and searches. Instead, we present a comparison of the PULpy predictions and PULDB on a well-known rumen bacterium, *Prevotella ruminicola* 23^25^ (assembly accession number GCA_000025925.1). First described by Bryant *et al* ^26^, *Prevotella* are capable of the fermentation of a range of complex and simple sugars.

According to PULDB, there are 24 PUL annotated on the *Prevotella ruminicola* 23 genome. On the same genome, our analysis using PULpy produces 22 predicted PUL. Some PUL predicted by one method overlap multiple PUL predicted by the other method, as PUL boundaries are difficult to define. A table showing how the predictions overlap is provided in Supplementary Table 2, and images displaying comparisons are available in Supplementary Figures 1 to 21 (created using ProGenExpress^27^). ALL PUL predicted by PULpy overlap with at least one of the PULDB annotations (i.e. there are no false positives). PULDB contains one PUL that PULpy misses (“Predicted PUL 18”) (i.e. there is one false negative).

Seven of the PULDB and PULpy predictions are identical, and these are listed in Table 1.

**Table 1.**
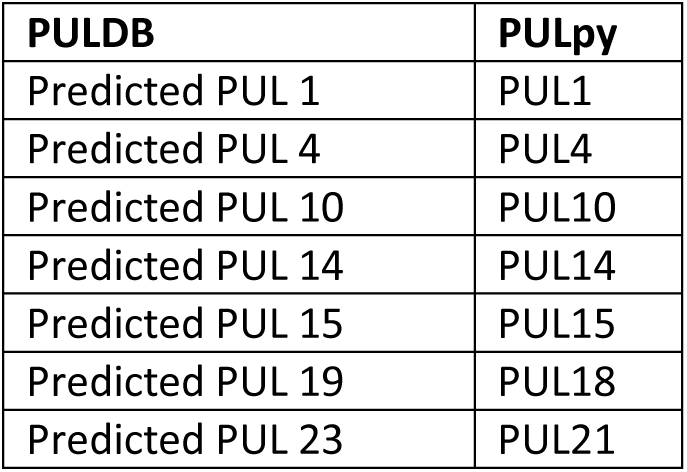
Identical PUL predictions from PULDB and PULpy

Many of the PUL comparisons are worthy of discussion. Supplementary Figure 1 shows PULDB’s “Predicted PUL 1” and PULpy’s PUL1, which are identical. Importantly, this PUL establishes the rule that it is valid to extend the PUL boundaries to include a CAZyme that is on the *opposite strand* to the *susCD* pair. This is important, as it appears this “rule” is not consistently followed within PULDB (though it is within PULpy).

Supplementary Figure 2 demonstrates a known weakness of PULpy. Here, “Predicted PUL 2” from PULDB has been extended to include an HTCS gene. HTCS genes are identified by PULDB as potential regulators^12^, but at present PULpy does not include these.

Supplementary Figure 3 compares “Predicted PUL 3” with PUL3. These are identical apart from the inclusion of two genes in “Predicted PUL 3”, one at either end, neither of which appear to have a function relevant to PUL. PULpy does not recognise them as CAZymes nor as *susCD*, therefore they are not included.

Supplementary Figure 5 shows a more complex example. Downstream of PUL5, PULpy has failed to include an HTCS gene for the reasons mentioned above. PUL5 also extends upstream to a glycosyl transferase gene not included in “Predicted PUL 5”. This gene is a CAZyme, is within the five-gene window, and none of the intergenic distances are greater than 500 bp. We are unsure why this is not included in the PULDB prediction. It *is* on the opposite strand to the *susCD* pair; however, as we have seen from “Predicted PUL 1”, this is allowed in some cases.

Supplementary Figure 6 contains PULDB’s “Predicted PUL 6”, “Predicted PUL 7”, and PULpy’s PUL6 and PUL7. A disagreement occurs over the membership of a glycoside hydrolase gene, which PULDB believes belongs to “Predicted PUL 7” whereas PULpy believes it belongs in PUL6. The large intergenic distance is key – the next closest coding sequence (CDS) to the disputed GH2 gene within “Predicted PUL 7” is 1170 bp away, greater than the 500 bp limit. The GH2 and CE12 gene (upstream) are separated by only 242 bp, so it is unclear why the two predictions in PULDB are not joined.

“Predicted PUL 8” and PUL8 are compared in Supplementary Figure 7. Here, the PULpy prediction has extended downstream to include an additional *susC* gene which PULDB has ignored. Whilst PULpy consistently follows the rules we set it, PULDB is less consistent – for example, “Predicted PUL 24” from PULDB shows a very similar example where the PUL *is* extended downstream to include a *susC* gene. It is impossible to extract from PULDB which rule is “correct”.

In Supplementary Figure 8, “Predicted PUL 9” includes a gene which is neither predicted as a CAZyme nor a *susCD* by PULpy, therefore it is ignored. Supplementary Figure 10 shows “Predicted PUL 11” containing a GH3 gene that is 1255 bp away from the nearest CDS. This limit is greater than the 500 bp imposed by PULpy, and therefore it is not included in PUL11. Supplementary Figure 11 shows another example where the PULpy prediction is extended to include a nearby *susC* gene, whereas the PULDB prediction is not.

Supplementary Figure 12 compares “Predicted PUL 13” to PUL13. Here the PULpy prediction has been extended to include a number of nearby CAZymes not included in the PULDB prediction. The intergenic distance between PRU_2185 and the next GH31 gene is only 255 bp, well within the 500 bp limit.

Supplementary Figure 15 is another example of PULDB including genes that are ignored by PULpy, essentially because PULpy only includes genes it can annotate as a CAZyme or *susCD*. Supplementary Figure 16 is similar, with the PULDB prediction extending one gene either side to include HTCS genes.

Supplementary Figure 17 contains the false negative result – here PULDB has “Predicted PUL 18” whereas PULpy shows nothing. The reason for the false negative is that PULpy fails to recognise protein ADE82049.1 as having a *susD*-like domain. The Pfam results for this protein show a split-match for Pfam entry PF12771.6 (“SusD-like_2”) against ADE82049.1, but neither of the split-matches individually pass PULpy’s filters. Having failed to identify a *susCD* pair in the region, PULpy cannot predict the rest of the PUL. The reason for the split-match is not clear – of the 488 positions within the HMM, 79 are not matched against ADE82049.1, resulting in the split-match.

Supplementary Figure 19 will take some explanation. There are three PULDB predictions (“Predicted PUL 20”, “Predicted PUL 21”, and “Predicted PUL 22”) and three PULpy predictions (PUL19 and PUL 20). Beginning at the left of the figure, we see an HTCS gene that PULpy does not cover. PUL19 largely overlaps “Predicted PUL 20”, but then extends downstream to include two glycosyl transferase genes that are not included in “Predicted PUL 20”. Having made that initial extension, PULpy is able to extend further to a glycoside hydrolase gene (five genes away), now over-lapping what PULDB annotates as “Predicted PUL 21”. That extension from the GT4 gene to the GH3 gene does not break any rules – the GH3 gene is within the five gene window, and none of the intergenic distances exceed 500 bp. PUL19 then finishes at a GH43_2 gene – the next genes downstream are a *susCD* pair, but these are greater than 500 bp away. However, “Predicted PUL 21”, which overlaps with the end of PUL19, *does extend* to include the very same *susCD* pair, despite the intergenic distance being 548 bp (greater than the 500 bp cut-off). Finally, PULpy’s PUL20 begins at this *susCD* pair and then extends to overlap with “Predicted PUL 22”. This is clearly an enormously complex region of the *Prevotella ruminicola* 23 genome in terms of the abundance of *susCD* pairs and CAZymes. It will always be very difficult to annotate PUL in this region accurately from genomic data alone. Clearly we should treat both PULDB and PULpy results in this region as predictions, to be tested with further experiments and data.

Finally, Supplementary Figure 21 contains “Predicted PUL 24” and PUL22. Here we see an important example from PULDB where a PUL is extended to include a single *susC* gene. This rule appears not to be adhered to throughout PULDB, but is allowed within PULpy. The reason PULpy does not include the additional *susC* gene is that PULpy does not recognise the gene as having any domains associated with *susC* – on this protein the Pfam results annotate no domains at all.

The PULpy predictions we present here are exactly that – predictions. As PULDB includes additional manual annotation, it makes sense to defer to PULDB where there is a disagreement. The advantage of PULpy over PULDB is that the PULpy predictions are open, reproducible, and extendible to newly sequenced organisms. Many of the PUL in PULDB are also predictions, and true PUL can only be defined with additional experimental data. We believe the PULpy predictions represent a useful first-pass annotation at PUL, and this release of the first fully open dataset and fully open tool for PUL prediction in *Bacteroidetes* genomes represents a useful advance.

## Acknowledgements

The Roslin Institute forms part of the Royal (Dick) School of Veterinary Studies, University of Edinburgh. This project was supported by the Biotechnology and Biological Sciences Research Council (BBSRC; BB/N016742/1, BB/N01720X/1), including institute strategic programme and national capability awards to The Roslin Institute (BBSRC: BB/P013759/1, BB/P013732/1, BB/J004235/1, BB/J004243/1); and by the Scottish Government as part of the 2016–2021 commission.

## Author contributions

Each author’s contribution to the work should be described briefly, on a separate line, in the Author Contributions section.

